# Vaginal Tissue Engineering via Gelatin-Elastin Fiber-Reinforced Hydrogels

**DOI:** 10.1101/2024.09.09.611932

**Authors:** Samantha G. Zambuto, Samyuktha S. Kolluru, Abir Hamdaoui, Annabella M. Mascot, Siobhan S. Sutcliffe, Jerry L. Lowder, Michelle L. Oyen

## Abstract

The vagina is a fibromuscular tube-shaped organ spanning from the hymenal ring to the cervix that plays critical roles in menstruation, pregnancy, and female sexual health. Vaginal tissue constituents, including cells and extracellular matrix components, contribute to tissue structure, function, and prevention of injury. However, much microstructural function remains unknown, including how the fiber-cell and cell-cell interactions influence macromechanical properties. A deeper understanding of these interactions will provide critical information needed to reduce and prevent vaginal injuries. Our objectives for this work herein are to first engineer a suite of biomaterials for vaginal tissue engineering and second to characterize the performance of these biomaterials in the vaginal microenvironment. We successfully created fiber-reinforced hydrogels of gelatin-elastin electrospun fibers infiltrated with gelatin methacryloyl hydrogels. These composites recapitulate vaginal material properties, including stiffness, and are compatible with the vaginal microenvironment: biocompatible with primary vaginal epithelial cells and in acidic conditions. This work significantly advances progress in vaginal tissue engineering by developing novel materials and developing a state-of-the-art tissue engineered vagina.

## 1. Introduction

### Background

Women, females, and other gender minorities have been previously understudied and excluded in basic and translational scientific research^1,2^. Unfortunately, as such, this has led to a dearth of sex-and gender-specific basic science information^3–6^. In recent years, women’s health research has seen explosive growth, with many more researchers challenging their fields^7,8^ to promote equitable research practices to propel inventions and innovations for women, females, and other gender minorities. Despite the rapid expansion of this field, many questions remain in this space that should receive attention.

One such question is how we can prevent and target gynecological injuries. Quality of life is severely negatively impacted by gynecologic conditions and obstetric injuries, including congenital malformations, childbirth injuries, bacterial vaginosis, vulvovaginal candidiasis, sexually transmitted infections, and the genitourinary syndrome of menopause. Unfortunately, there are limited treatment options for many of these conditions, and innovating in this space is challenging due to a dearth of basic science data related to gynecological function. The generation of basic science data pertaining to female gynecology has strong potential to fuel innovations in this space by providing crucial basic science information on gynecological anatomy and physiology, especially related to the vagina.

### The Vagina

The vagina is a fibromuscular tube-shaped organ spanning from the hymenal ring to the cervix that plays critical roles in menstruation, pregnancy, and female sexual health^9^. The vagina is approximately 6-10 cm in length, with wall length and diameter varying from posterior to anterior direction and cervix to introitus^10–12^. Vaginal tissue is composed of 4 distinct layers (**Fig. 1**): epithelium, lamina propria, muscularis, and adventitia^10,13,14^. In women of reproductive age, the vaginal epithelium contains approximately 28 layers of non-keratinized stratified squamous epithelium that are entirely replaced every 96 hours^10^. The lamina propria is vascularized and contains fibroblasts, immune cells (e.g., Langerhans cells, macrophages, dendritic cells, lymphocytes), and extracellular matrix (ECM) components–primarily collagen, which gives the vagina high tensile strength, and elastin, which enables the tissue to withstand deformation, especially in childbirth^10,13^. The muscularis contains smooth muscle cells organized into bundles oriented in circular and longitudinal directions^15^. The adventitia connects the vaginal tissue to surrounding structures. Another important vaginal component is the vaginal microbiome. It is composed primarily of lactobacilli, which play an essential role in maintaining the low pH of the vaginal microenvironment and protecting the tissue against various infections, and reducing susceptibility to certain sexually transmitted diseases ^16–18^.

**Figure 1.**
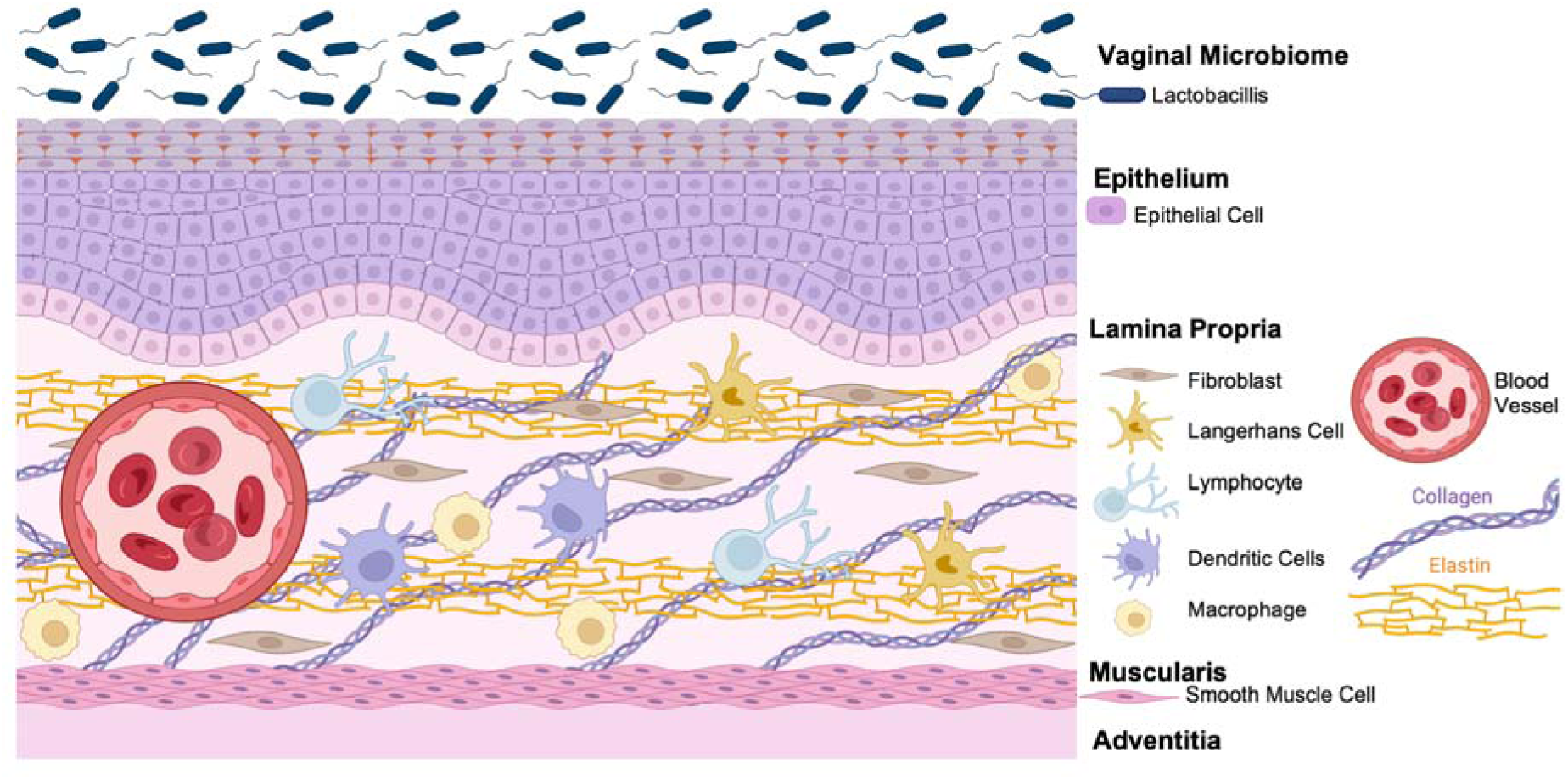
The vaginal tissue microenvironment

### Models of the Vagina

Animal models, including rodents, rabbits, sheep, minipig, and non-human primates, have been used to investigate the vagina and generate information vaginal functions^10,19^; however, many animal models are not ideal for many reasons. For example, the human bipedal position causes a unique pelvic floor orientation in humans, and the pelvic floor muscles support the pelvic organs by resisting abdominal pressure and gravity^20^. In contrast, most animals, including widely used rodent animal models, are quadrupeds. Their pelvic floor muscles also control their tails, thus shifting support of the pelvic viscera to the abdominal wall^20^. Additional differences include hormonal differences between humans and animals after birth and a larger fetal head-to-vaginal canal size ratio in humans than in animals^20^. Non-human primate animal models are advantageous in that primate reproduction and the primate vaginal microbiome is similar to human reproduction and the human vaginal microbiome^10,19^; however, ethical concerns and high costs limit the use of primates for research purposes.

A benefit to *in vitro* model systems for studying the vagina is that direct mechanistic effects can be explored in highly tunable, high throughput systems. In a recent review by McCraken et al., the authors highlight important considerations for creating nonclinical models of the vagina^10^. Notably, the authors point out that many existing *in vitro* vaginal models cannot recapitulate important biomechanical and cellular vaginal features, including Valsalva pressures to the vagina, vaginal tensile strength, cellular compositions, and permeability properties^10^. Notably, many of these features that need to be added can be simulated using tissue engineered model systems.

### Hydrogels as Conduits for Vaginal Tissue Engineering

Hydrogels are excellent candidates for vaginal tissue engineering due to their hydrophilicity, biocompatibility, and ease of use; however, hydrogels are soft and compliant, which is not conducive to mimicking specific tissues like the vagina^21^. Hydrogel-fiber composites are amenable to tissue engineering constructs because they share the properties of hydrogels but with increased load-bearing capacity from fibers^21^. Fibers can be generated using electrospinning, a method in which polymer fibers are produced by applying a large voltage between the polymer solution and grounded target^21,22^. A hydrogel-fiber composite is highly attractive for vaginal tissue engineering because biocompatibility, biophysical characteristics (e.g., fiber alignment), and higher biomechanical strength, toughness, and stiffness can be generated with these fiber-reinforced composites. We have well-established techniques for creating fiber-reinforced composites using a variety of biomaterials^21,23–27^.

Nonclinical engineered model systems of the human vagina are needed to investigate vaginal biomechanics while replicating relevant features of the human vagina, including cellular composition and architecture, geometry, and hormone levels. The long-term goal of this project is to establish a tissue engineered vagina model. Our objectives for this study were to engineer a suite of biomaterials for vaginal tissue engineering and characterize the performance of these biomaterials in the vaginal microenvironment. We created fiber-reinforced hydrogels consisting of gelatin-elastin electrospun fibers infiltrated with viscoelastic gelatin methacryloyl (GelMA) hydrogels. These biomaterials and resultant tissue engineered vagina model can be used as tools to continue building tissue engineered vaginal models of increasing complexity and can provide engineered platforms to define vaginal biomechanics and generate crucial basics science data to target and prevent vaginal injury.

## 2. Materials and Methods

### 2.1. Electrospun fiber synthesis and characterization

#### 2.1.1. Production of Electrospun Fibers

Gelatin (Sigma Aldrich G2500), soluble elastin (Elastin Products Company Inc., ES12), and gelatin and elastin blends in varying ratios (1:2, 1:1, 2:1) were dissolved until homogeneous in a 10 wt% solution containing acetic acid (90% aqueous solution; Sigma Aldrich A6283), citric acid (15 wt% based on gelatin weight; Sigma Aldrich 77-92-9), and sodium hypophosphite (7.5 wt%; Sigma Aldrich 10039-56-2). Fibers were spun for 6 hours using the following electrospinning parameters: applied voltage of 10 kV, 0.3 mL/hr rate of extrusion, syringe diameter of 20 mm, 2-minute cleaning frequency, 50 mm spinneret width, 12 mm/second spinneret speed, and approximately 10 cm distance between needle tip and grounded collector. Citric acid and sodium hypophosphite were crosslinking catalysts, and the resultant electrospun mat was placed in a 150°C oven for 4 hours for crosslinking via dehydration^28^.

#### 2.1.2 Dynamic Viscosity Measurements

Electrospinning solutions were added to a Cannon-Fenske Viscometer. The time for the solution to travel through the viscometer was measured using a stopwatch. These data were subsequently analyzed via a custom MATLAB code to determine kinematic viscosity using **Equation 1** and dynamic viscosity using **Equation 2**. The custom MATLAB code can be found in the **Supplemental Information**.

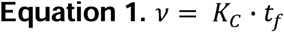

Where *v* is kinematic viscosity calculated by the product of the viscometer constant (*K_C_*) and measured flow time (*t_f_*).

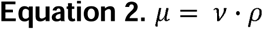

Where *µ* is dynamic viscosity calculated by the product of the kinematic viscosity and density (*p*).

#### 2.1.3. Attenuated Total Reflectance-Fourier Transform Infrared Spectroscopy

Fibers and powdered stock samples were analyzed via attenuated total reflectance-Fourier transform infrared spectroscopy (ATR-FTIR) using a Bruker Alpha II FTIR outfitted with a zinc selenide crystal (spectral range 20,000 – 500 cm^-1^). A background scan was performed before sample measurement, and spectra were averaged over 24 scans, recorded from 600 – 4000 cm^-1^ with a resolution of 4 cm^-1^.

#### 2.1.4. Environmental Scanning Electron Microscope Imaging and Fiber Size Analysis

Ten regions of interest were imaged for crosslinked and uncrosslinked 2:1 gelatin:elastin fibers using an Environmental Scanning Electron Microscope Thermofisher Quattro S ESEM. The fibers were attached to the pin stub using carbon tape. Images were taken at 20,000X magnification, 2.0 kV, and 3.0 spot size. Fiber diameters were measured on ImageJ using the Measure tool. Individual fiber diameters were measured from both sides of the image while maintaining a consistent orientation perpendicular to the original direction of the fibers. At least thirty fibers were selected in each image at approximately the same imaging plane.

### 2.2. Gelatin methacryloyl (GelMA) synthesis and characterization

#### 2.2.1. GelMA Synthesis

Gelatin methacryloyl (GelMA) of approximately 50% degree of modification was synthesized as described previously using the one pot method ^29^. High bloom (300 g) Porcine gelatin type A (Sigma Aldrich G2500) was dissolved 10% w/v in carbonate buffer (ThermoFisher 28382) at 50°C and methacrylic anhydride (40 μL/g gelatin; Sigma Aldrich 276685) was added dropwise to the solution while stirring. The reaction was allowed to proceed for 1h and was subsequently stopped with the addition of deionized water (40 mL/g gelatin). Once the pH of the solution was adjusted to 6 – 7, the resultant solution was dialyzed against deionized water for seven days (12 -14 kDa molecular weight cut off; Fisher Scientific 21-152-8) and lyophilized.

### 2.4. Fiber-reinforced hydrogel composite fabrication and characterization

#### 2.4.1. Fiber-reinforced Hydrogel Fabrication

A 5 wt% (weight percent) hydrogel prepolymer solution was prepared by dissolving lyophilized GelMA in phosphate-buffered saline (PBS) at 37°C. Once the solution was dissolved, 0.1% w/v lithium acylphosphinate (LAP; Sigma Aldrich 900889-1G) photoinitiator was added to the solution and thoroughly mixed. The 2:1 gelatin:elastin fibers (10 mm diameter) were punched out with a biopsy punch and placed into a custom Teflon mold (10 mm diam., 1 mm height). A dipcoat method was used to create composites. Briefly, prepolymer solution was pipetted (100 μL) onto the fibers, and the resultant composite was then moved to another well on the mold. Composites were polymerized for 5 minutes under UV light (λ = 365 nm; Spectroline EA-160), carefully removed from the molds using tweezers and placed in 24 well plates filled with PBS. After critical point drying, ESEM images were taken of the composites, homogenous hydrogels, and fibers. All relevant instrument settings are reported in each shown ESEM image.

#### 2.4.2. Uniaxial Tensile Testing

Fiber mats were cut into 30x10 mm rectangular specimens, and the edge of the mat was discarded to ensure no anomalies arising from the spinning process were included in the tested specimens. Next, 100 μL prepolymer solution was pipetted onto the mat for infiltration, and then the mat was moved to a new well to create composites of similar stiffness as the 10 mm samples using a dip-coat method. Thickness was measured using digital calipers. The fiber mats/composites were cut into dog bone shapes, with a 5 mm width at the narrowest region. This ensured that the stress was concentrated at the center of the sample. Uniaxial tensile testing was performed on an Instron (Model 68SC-1) mechanical testing system with a 50N load cell and a tear rate of 0.5 mm·s^-1^.

The data were used to calculate elastic modulus (*E*), ultimate tensile strength (σ_UTS_), strain energy density (*U*), and failure strain (*e*) with *n* = 8 samples per condition. Elastic modulus was determined from the slope of the linear regime of the stress-strain curve. The maximum stress value during the tensile test is the ultimate tensile strength. Failure strain is the maximum strain value. Strain energy density is calculated from the area under the stress-strain curve and is defined as the energy dissipated per unit volume. All calculations were performed using a custom MATLAB code (code available on GitHub at: https://github.com/sskolluru/Tear-and-tensile-testing-analysis-codes).

Trouser tear testing followed the same protocol as above, except that instead of a dog bone shape, the sample was made with a 10 mm notch in a rectangular mat that resembled a pair of trousers. The data were used to calculate tear toughness, with *n* = 8 samples per condition.

#### 2.4.3. Mass Swelling Ratio

Composites were created and submerged in PBS for 24 hours to ensure they reached equilibrium swelling. After weighing the swollen samples, the samples were lyophilized until dry and weighed once more. Mass swelling ratio (*q*), a proxy for crosslinking density, was calculated using **Equation 3**

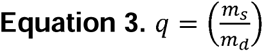

by comparing the swollen composite weight (*m_s_*) and lyophilized (dry) composite weight (*m_d_*).

#### 2.4.4. Degradation Assay

Vaginal fluid simulant was prepared following the protocol established by Rastogi et al.^30^. The 10X vaginal fluid simulant buffer was made by combining 5.25 g of sodium chloride (NaCl; Fisher Scientific S642-500) with 2.02 g of sodium lactate (diluted to 50% w/w; Fisher Scientific AA41529AK), adding 790 μL of acetic acid (Sigma Aldrich A6283), and bringing the volume to 100 mL using distilled water. The pH was measured using pH strips and determined to be 4.2. The final concentrated vaginal fluid simulant was then stored at 4°C and diluted 10X with distilled water before use. Composites were created and submerged in PBS for 24 hours to reach equilibrium swelling. After 24 hours, samples were transferred into well plates filled with vaginal fluid simulant solution, and their mass was measured at different time points. The degree of degradation (*α*) was calculated using **Equation 4**.

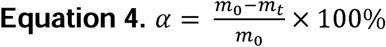

Where *m_0_* is the initial sample mass after equilibrium swelling and *m_t_* is the sample mass at a specific time.

### 2.5. Primary Vaginal Epithelial Cell Culture

#### 2.5.1. General Cell Culture

Primary human vaginal epithelial cells (LifeLine Cell Technology FC-0083, Lot 02841) were cultured as per the manufacturer’s instructions in phenol red-free ReproLife^TM^ Complete Medium (LifeLine Cell Technology FC-0078). Cells were passaged only once after receipt. The vendor reports these are female cells from a 28-year-old Caucasian woman. Cell-laden composites were created by adding 500,000 cells/mL to the prepolymer solution and subsequently polymerizing the composites under UV light, as described in section 2.4.1. Cell-laden composites were cultured in 24 well plates containing 1 mL ReproLife^TM^ Complete Medium per well. All cells were cultured at 37℃ in 5% CO_2_ incubators.

#### 2.5.2. Live/Dead Staining

After 24 hours of culture, cell-laden composites were stained with a LIVE/DEAD Viability/Cytotoxicity Kit (Invitrogen L3224) for 20 minutes in the incubator. Following incubation, a 5–10-minute PBS wash step was performed, and washed samples were stored in PBS for imaging. Samples were imaged on a Keyence All-in-One Fluorescence Microscope BZ-X series (KEYENCE Corporation of America).

#### 2.5.3. Nascent Protein Staining

We conducted nascent protein staining as described previously^31,32^ using the methods developed by Loebel et al.^33,34^. Briefly, after fabrication, cell-laden composites were incubated in Hank’s Balanced Salt Solution (HBSS) for at least 30 minutes to deplete endogenous methionine. Next, composites were incubated in 100 μM of the methionine analog azidohomoalanine Click-iT AHA (Invitrogen C10102) in HBSS for 1 hour in the incubator. Two HBSS washes were performed, and then the composites were incubated in 30 μM AFDye 488 DBCO (Click Chemistry Tools 1278-1) for 40 minutes in the incubator. Composites were washed thrice with PBS following incubation and then fixed for 15 minutes in formalin. After fixation, composites were stained using 1:1000 CellMask^TM^ Deep Red Plasma Membrane Stain (Invitrogen, C10046) for 40 minutes at 37°C. Three PBS washes were performed, and then the composites were stained with 1:2000 Hoechst at room temperature for 30 minutes. Then, they were washed once following incubation. Composites were stored in PBS until imaging on the Keyence All-in-One Fluorescence Microscope BZ-X series (KEYENCE Corporation of America).

#### 2.5.4. ESEM Imaging

ESEM images of cell-laden composites were obtained following fixation with formalin and glutaraldehyde, freezing in liquid nitrogen, fracturing, critical point drying, and mounting on pin stubs. Samples were mounted and coated with 6 nm of carbon followed by 6 nm iridium and were stored in a vacuum desiccator until imaged on a Quattro S Environmental Scanning Electron Microscope (Thermo Scientific). Instrument settings are reported on each ESEM image.

### 2.3. Statistical Analysis

Descriptive statistics are reported in the results as mean ± standard deviation unless otherwise noted. The total sample number (*n*) for each experiment is reported in the figure caption. Statistical analyses were chosen after testing the assumptions for normal (Shapiro-Wilk test) and homoscedastic (Bartlett’s test) data. The statistical test selected is reported in the figure caption. Significance was set to p < 0.05. All quantitative plots were created using Prism 9 (GraphPad), and statistical analyses were performed using Prism 9 (GraphPad) or R (R-Project.org).

## 3. Results

### 3.1. Production and Characterization of Electrospun Gelatin-Elastin Meshes

Electrospinning 100% elastin resulted in significant sample beading (Fig. 2A). We, therefore, added gelatin as a copolymer to the solution. We spun gelatin:elastin fiber meshes in 3 ratios (1:1, 1:2, 2:1) and determined that the 2:1 gelatin:elastin ratio resulted in consistent fibers (**Fig 2A**). The 2:1 gelatin:elastin blend resulted in a viscosity (10.58 ± 4.71 cP) that was not statistically significantly different from the 100% gelatin solution (9.05 ± 0.08 cP). We determined that the addition of gelatin as a co-polymer increased solution viscosity and the increased viscosity is what allowed the fibers to be spun (**Fig. 2B**). The low viscosity of 100% elastin and the 1:2 and 1:1 gelatin:elastin blends (0.70 ± 0.01 cP, 1.14 ± 0.47 cP, 2.33 ± 0.23 cP, respectively) contributed to sample beading when electrospun (**Fig. 2B**).

**Figure 2.**
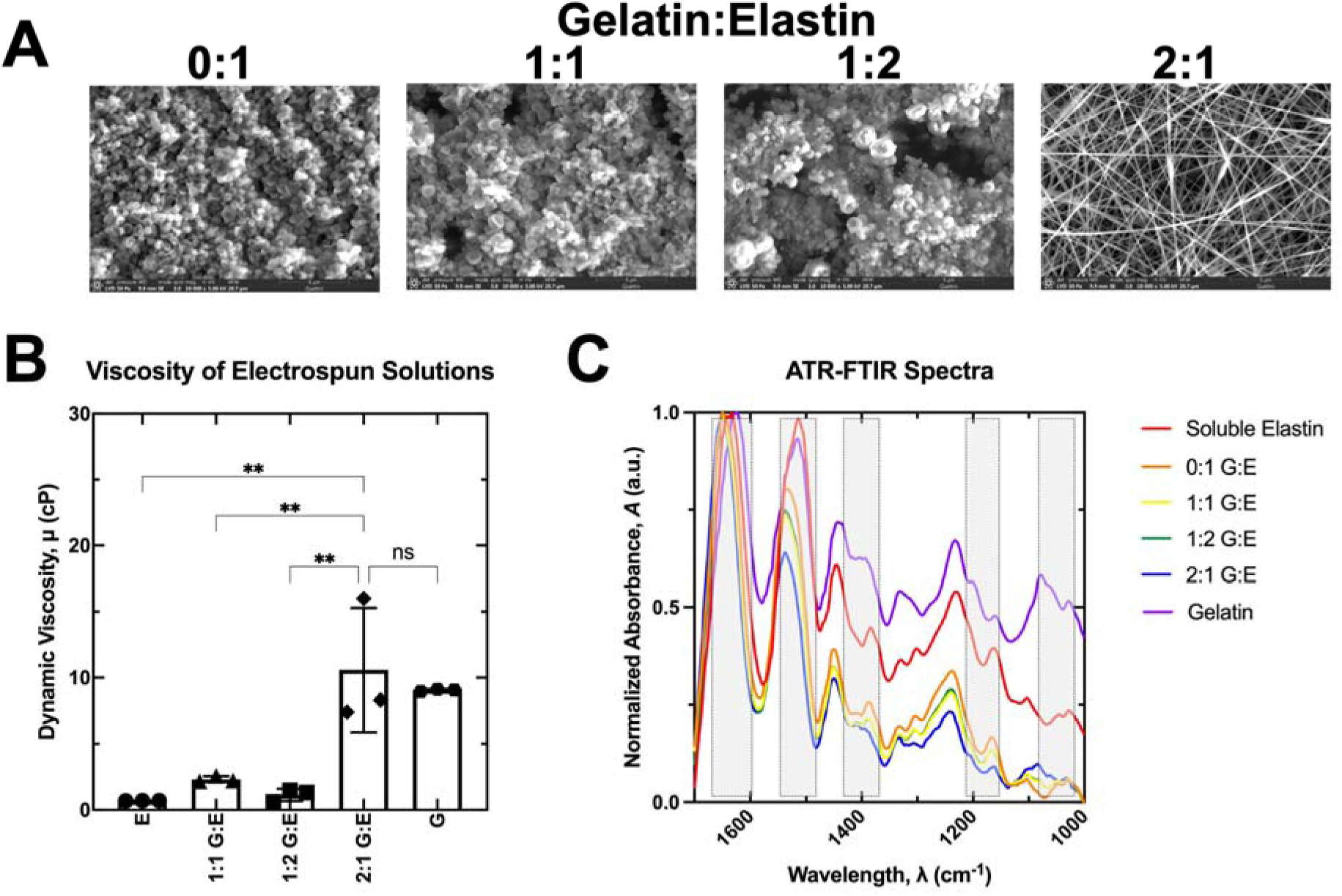
Electrospun gelatin:elastin fiber blends. **A.** Representative ESEM images of 0:1, 1:1, 1:2, and 2:1 gelatin:elastin fibers. **B.** Dynamic viscosity of electrospinning solutions. Statistical analysis via ordinary one-way ANOVA and Tukey posthoc analysis. Significance defined as p < 0.05 where ns (not significant) is p > 0.05 and ** is p < 0.01. Multiple comparisons are shown for comparisons between the 2:1 ratio and other ratios only. **C.** ATR-FTIR spectra for gelatin, elastin, and fibers. Truncated data from 1000-1700 cm^-1^ are normalized. Shaded regions represented regions of interest with differences in peaks between gelatin, elastin, and fibers. E: Elastin. G: Gelatin.

ATR-FTIR spectra analysis demonstrated the incorporation of elastin and gelatin into the fibers (**Fig. 2C**). We observed characteristic elastin peaks around 1600 cm^-1^ associated with amide I C=O stretching vibrations,1500 cm^-1^ associated amide II N-C=O stretching vibrations, and 1200 cm^-1^ associated with amide III^35–37^. We also observed characteristic gelatin peaks around 1600 cm^-1^ associated with amide I C=O stretching, 1500 cm^-1^ associated with N-H bending coupled to C-H stretching, and 1200 cm^-1^ associated with amide III C-N stretching and N-H bending^38^. We observed peaks associated with stock gelatin and stock elastin in the gelatin:elastin fiber blends in five regions (highlighted; **Fig. 2C**).

Crosslinked fibers had an average diameter of 101.9 ± 3.6 nm (range: 95.0 – 106.8 nm), whereas uncrosslinked fibers had an average diameter of 138.2 ± 19.0 nm (range: 112.9 – 164.1 nm) (**Fig. 3A,B**).

**Figure 3.**
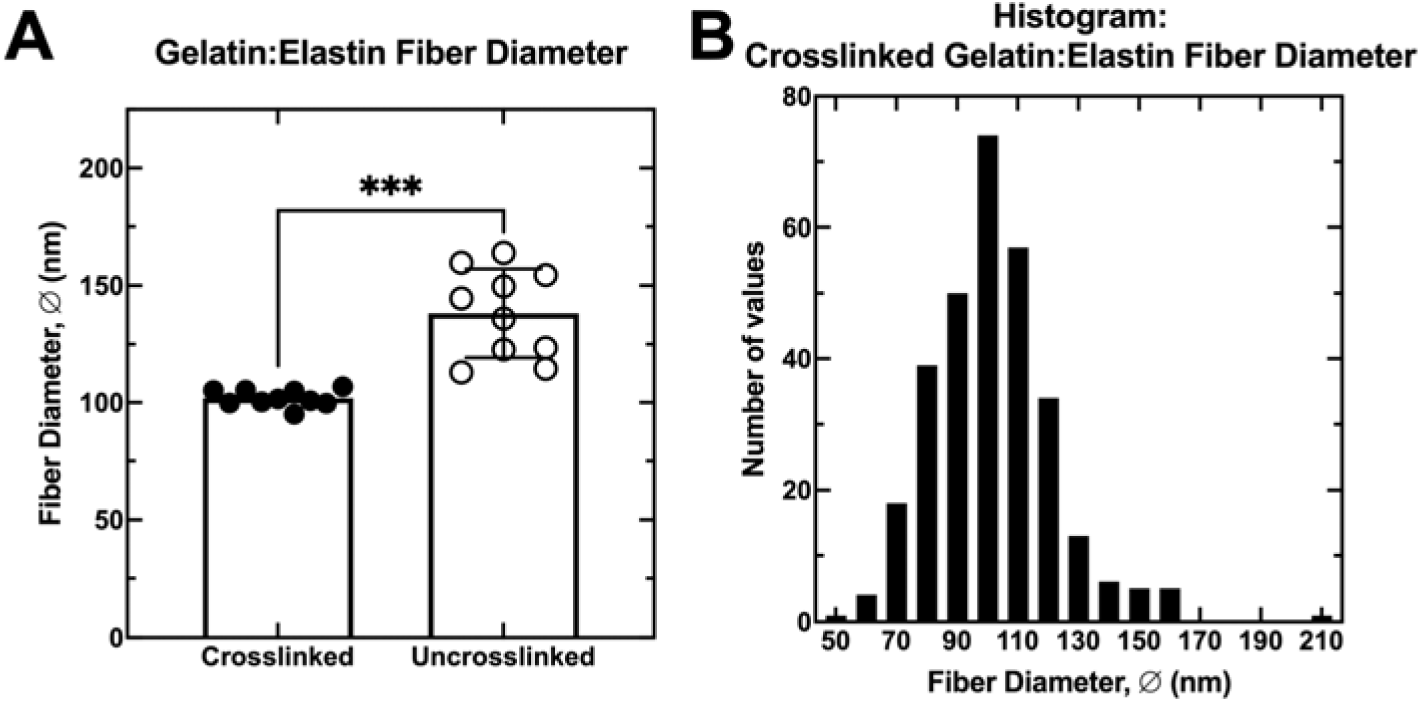
Electrospun fiber 2:1 gelatin:elastin fiber blend diameters. **A.** Fiber diameter mean ± standard deviation for crosslinked and uncrosslinked fibers. Statistical analysis via Welch’s t-test with significance set to p < 0.05. *** is p < 0.001. **B.** Histogram of crosslinked fiber diameters.

### 3.2. Material and Mechanical Properties of Fiber-Reinforced Composites

Fiber-reinforced composites were fabricated using a dip-coat method that ensured the fibers were infiltrated with hydrogel solution (**Fig. 4**). We then characterized the material and mechanical properties of the resultant composites. Material and mechanical properties of the composites are reported in **Table 1**, including work to failure, elastic modulus, strain energy density, ultimate tensile strength, failure strain, and fracture toughness. The mass swelling ratio of the composites was determined to be 21.3 ± 3.9 mg/mg. Graphs with individual data points for tear testing results and mass swelling can be found in supplemental **Figures S1** and **S2**.

**Figure 4.**
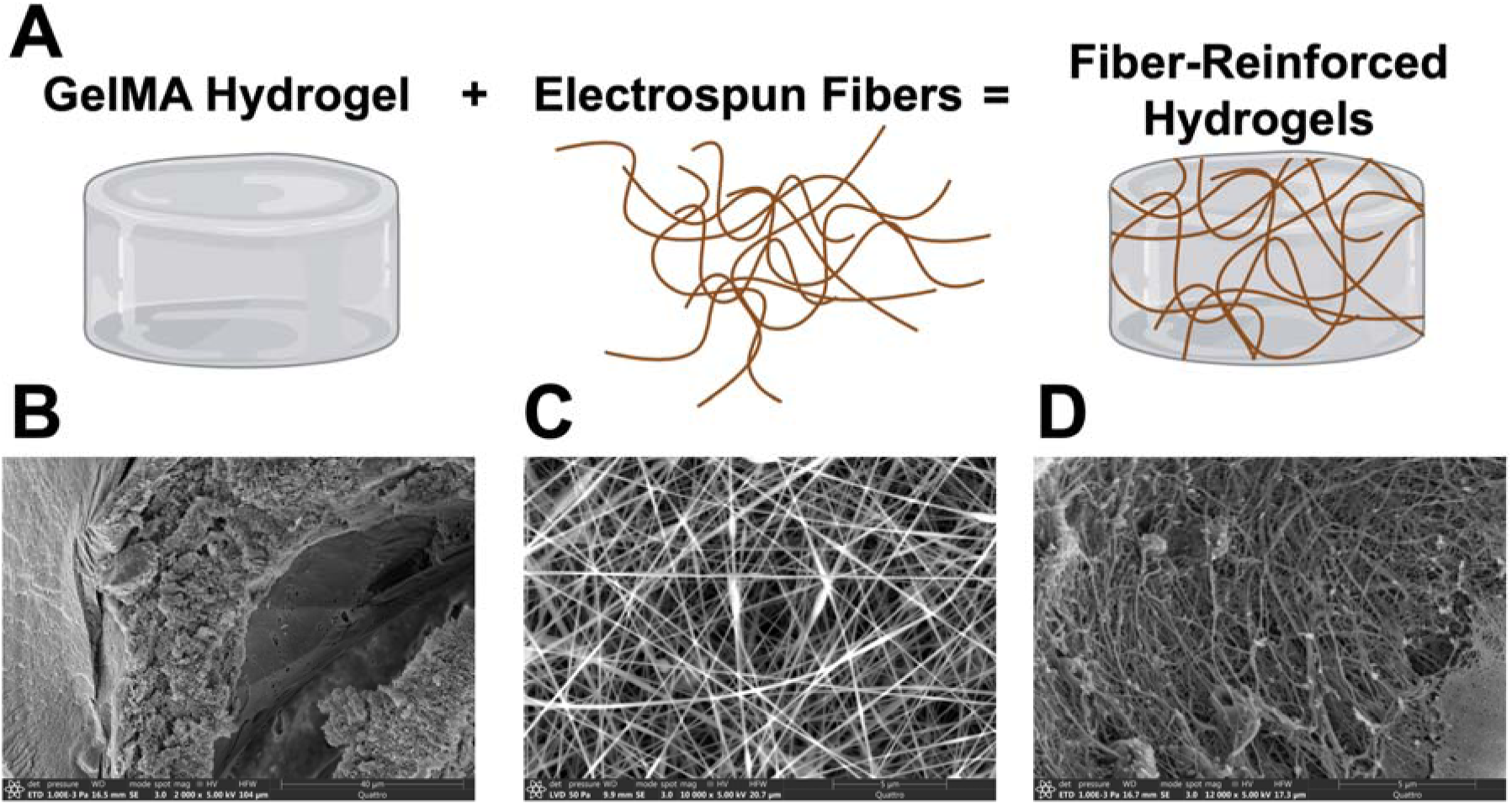
Fiber-reinforced hydrogel composites. **A.** Schematic of hydrogels, fibers, and fiber-reinforced composites. Representative ESEM images of **B.** a homogenous GelMA hydrogel, **C.** 2:1 gelatin:elastin electrospun fibers, and **D.** a fiber-reinforced composite.

**Table 1.** Material and Mechanical Properties of Fiber-Reinforced Composites.

| Property | Value (mean ± standard deviation) |
| --- | --- |
| Work to Failure ($W_f$) | $7.87 \pm 4.62$ kJ |
| Elastic Modulus ($E$) | $0.053 \pm 0.026$ MPa |
| Strain Energy Density ($U$) | $1.12 \times 10^{11} \pm 7.10 \times 10^{10}$ J/m<sup>3</sup> |
| Ultimate Tensile Strength ($\sigma_{UTS}$) | $0.017 \pm 0.0052$ MPa |
| Failure Strain ($\epsilon$) | $46.52 \pm 23.78$ % |
| Fracture Toughness ($T$) | $46.11 \pm 26.51$ J/m<sup>2</sup> |

### 3.3. Characterization of Electrospun Gelatin-Elastin Meshes in the Vaginal Microenvironment

Next, we sought to assess the performance of the resultant fiber-reinforced composites in the vaginal microenvironment. The vaginal microenvironment has a low pH, so we created a vaginal fluid simulant from protocols in the literature and quantified the degree of degradation of the composites, fibers, and homogenous hydrogels over time to assess their stability in low-pH environments (**Fig. 5**). Compared to homogenous hydrogels and fibers alone, the composites had the least degree of degradation and were the most stable over time. The fibers alone had the highest degree of degradation.

**Figure 5.**
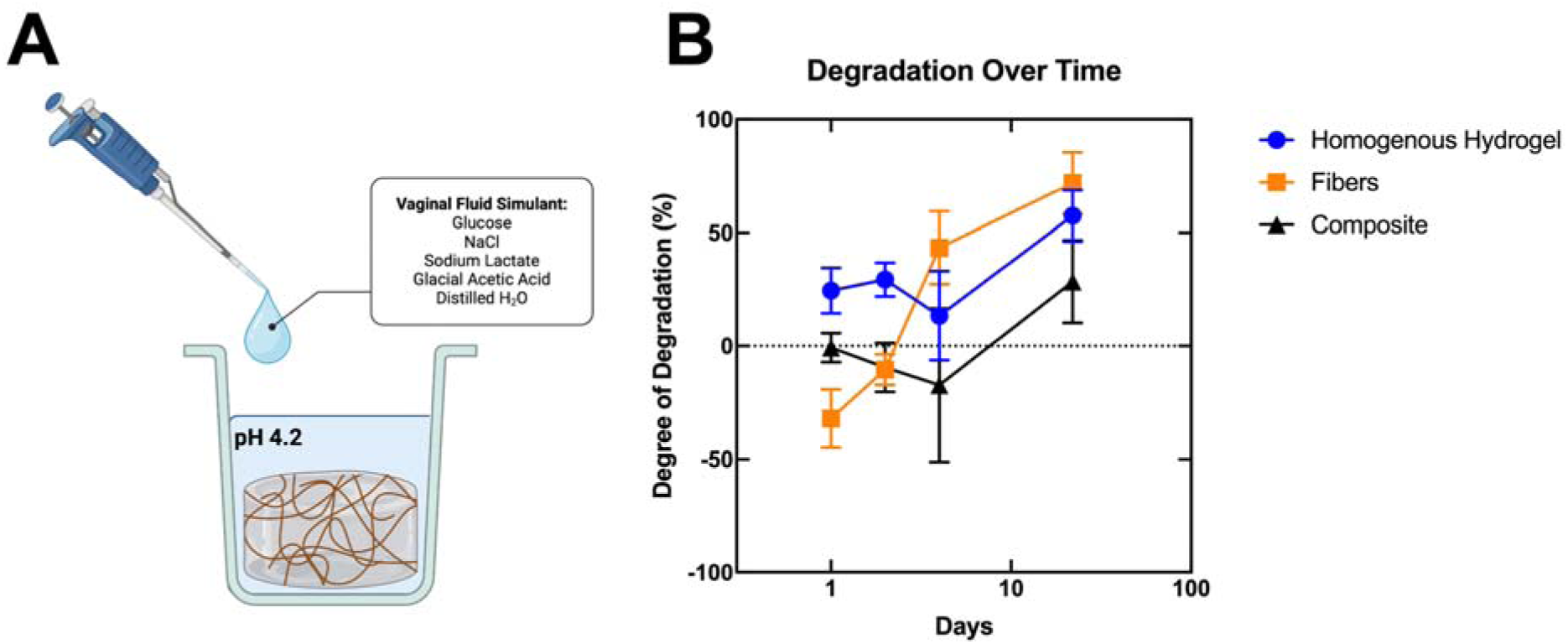
Degradation of composites in vaginal fluid simulant over time. **A.** Experimental schematic. **B.** Degradation over time for homogenous hydrogels, fibers, and composites.

We then created cell-laden composites by encapsulating primary human vaginal epithelial cells within the composites to assess biocompatibility and bioactivity of cells within the constructs (**Fig. 6A**). We observed high cell viability in the constructs as demonstrated through live/dead staining (**Fig. 6B**). We also demonstrated that the encapsulated vaginal cells were able to deposit nascent proteins within the composites (**Fig. 6C**), indicating that the cells are interacting with the composite. Finally, ESEM imaging revealed the incorporation of the cells within the composites and cell spreading (**Fig. 6D,E**).

**Figure 6.**
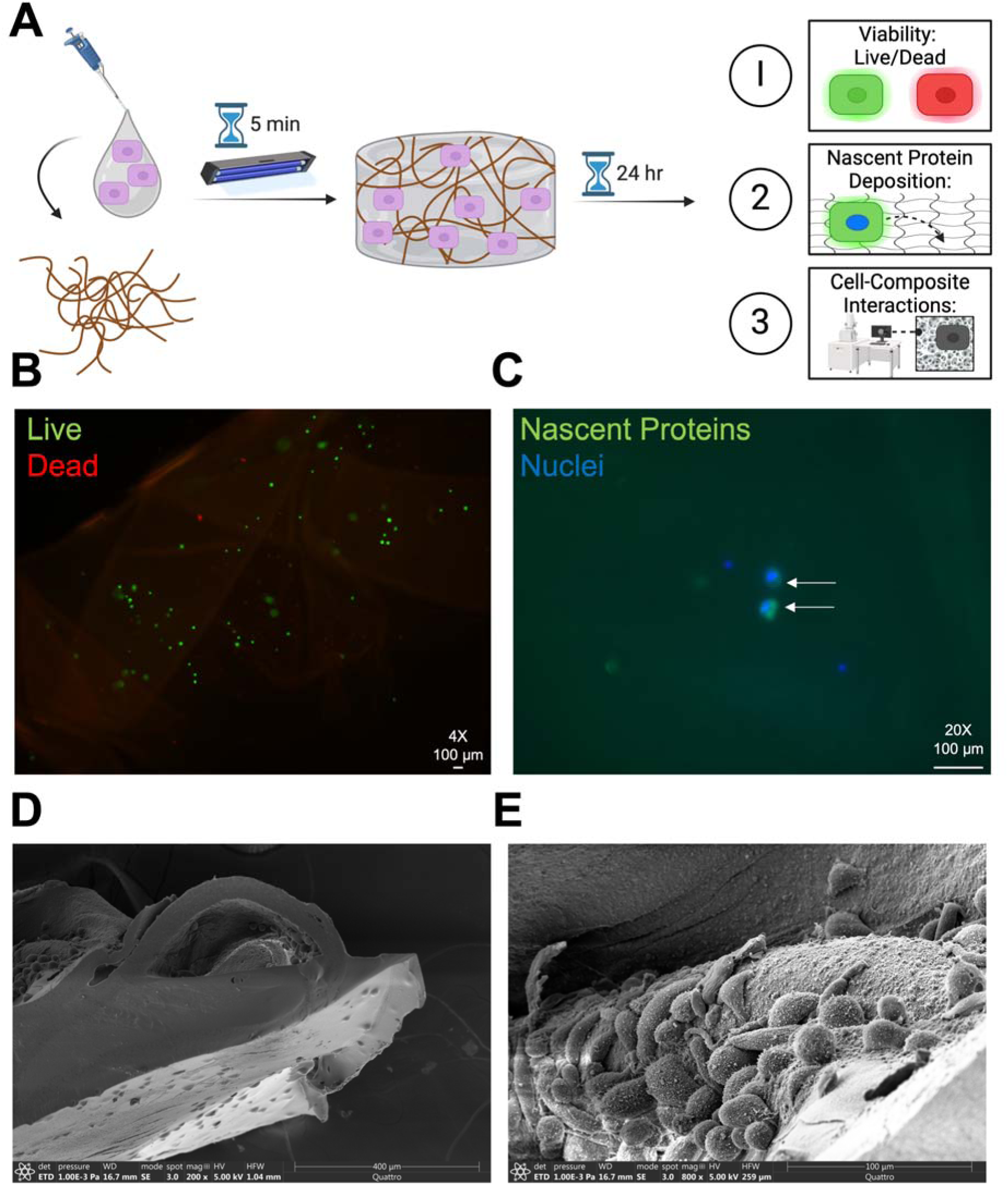
Vaginal cell interactions with fiber-reinforced composites. **A.** Experimental schematic. **B.** Live/Dead staining at 24h of culture. **C.** Nascent protein staining. White arrows: cells that deposited nascent proteins. **D.** and **E.** ESEM imaging of cell-laden composites at two magnifications.

Finally, we compared the elastic modulus of the composites to the elastic moduli of native vaginal tissue and homogenous GelMA hydrogels from two different studies (**Fig. 7**). The fiber-reinforced composites are two orders of magnitude higher than homogenous hydrogels and two orders of magnitude lower than vaginal tissue. Nonetheless, fiber-reinforced composites have an elastic modulus which more closely aligns with the elastic modulus of vaginal tissue compared to homogenous hydrogels.

**Figure 7.**
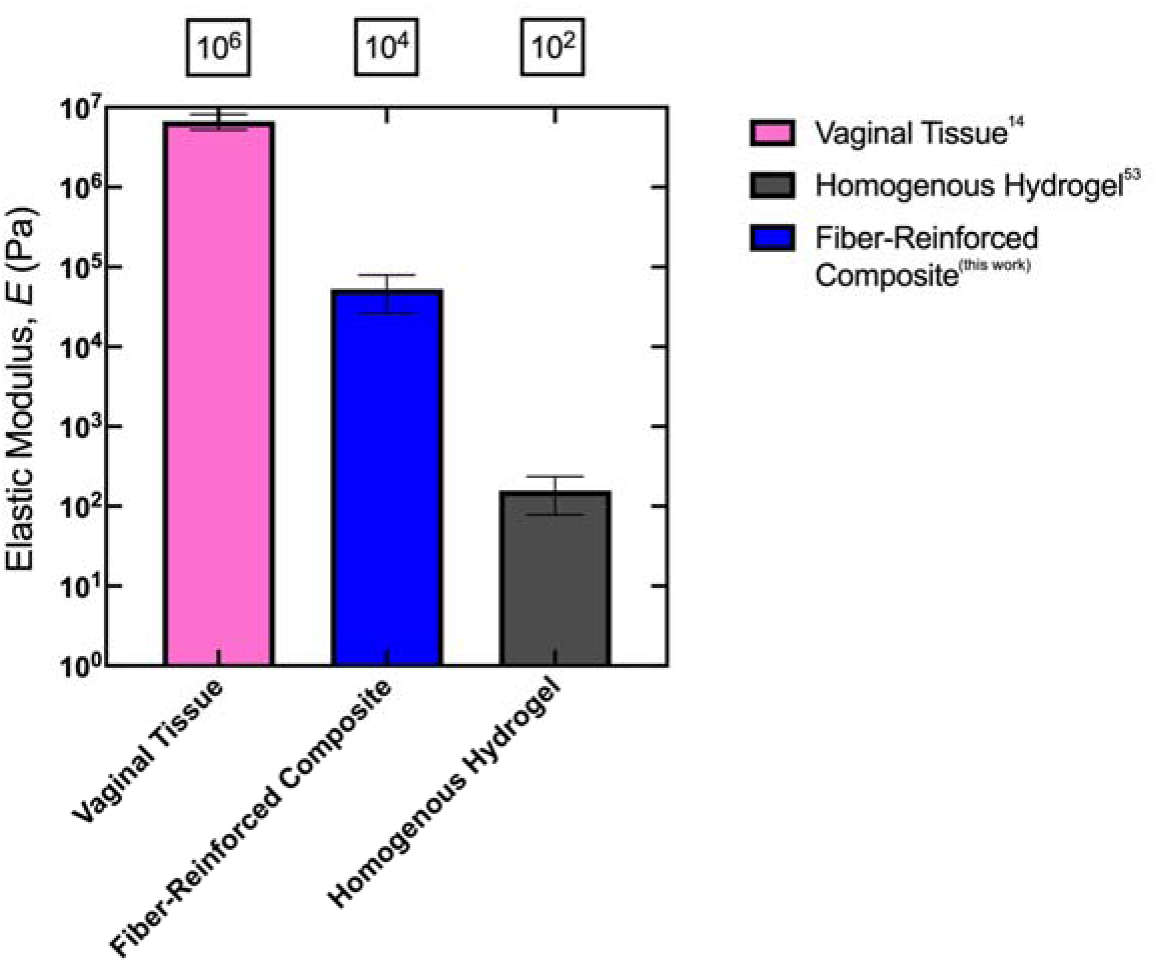
Comparison of elastic moduli of vaginal tissue, fiber-reinforced composites, and homogenous hydrogels. Vaginal tissue elastic modulus value from Gimenez et al., 2023^14^. Fiber-reinforced composite modulus from this work. Homogenous hydrogel modulus value from Zambuto et al., 2024^50^.

## 4. Discussion

Our long-term goal is to establish a tissue engineered vagina model to use for basic science studies of vaginal biomechanics. Our objectives for this work herein were to engineer a suite of biomaterials for vaginal tissue engineering and characterize the performance of these biomaterials in the vaginal microenvironment. Following our predefined engineering criteria, we successfully created fiber-reinforced hydrogels consisting of gelatin-elastin electrospun fiber blends infiltrated with gelatin methacryloyl (GelMA) hydrogels. We characterized the material and mechanical properties of these composites and tested their performance in the vaginal microenvironment.

For the first-generation composites, we chose the following engineering criteria as benchmarks: stiffness in the megapascal regime, viscoelasticity, vaginal epithelial cells as cellular constituents, and fiber components of collagen and elastin. These benchmarks were selected from our predefined engineering criteria (**Fig. 8**). Because interactions between collagen and elastin fibers play a crucial role in vaginal biomechanics, our model includes both types of fibers to recapitulate the native tissue structure^9^. We successfully achieved all the chosen benchmarks. We used gelatin-elastin fiber blends infiltrated with viscoelastic GelMA hydrogels. We determined that the fiber-reinforced composites had an elastic modulus closer to that of native vaginal tissue compared to homogeneous hydrogels that fell within the megapascal regime. We also determined that they are stable at a low pH like that of the *in vivo* vagina, and they maintain high cell viability, cell bioactivity, and cell-material interactions when laden with primary human vaginal epithelial cells.

**Figure 8.**
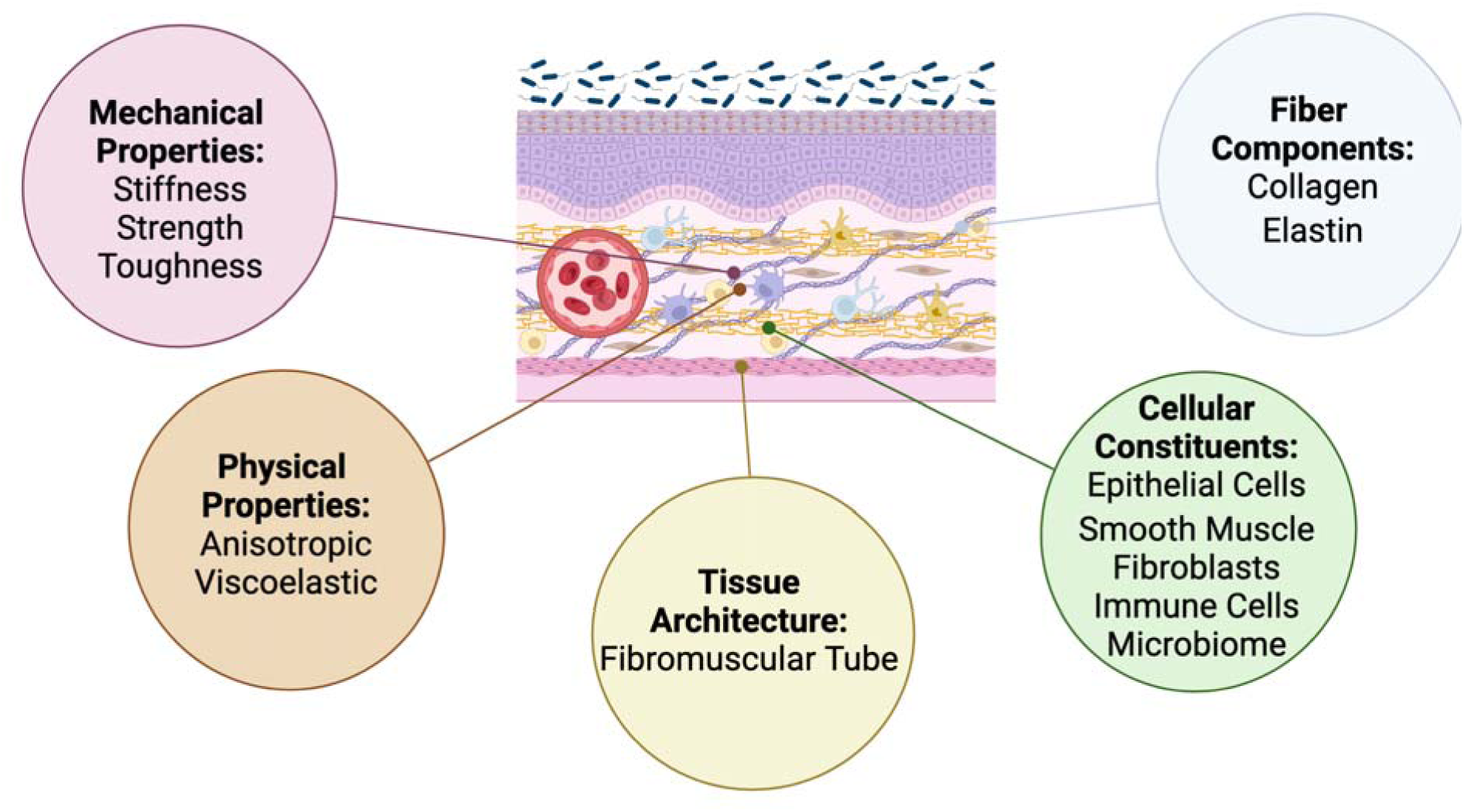
Engineering criteria for the design of a tissue engineered vagina.

The next-generation composites will focus on the following benchmarks (**Fig. 8**): increased composite strength and toughness, anisotropy, tube-shaped architecture, and vaginal smooth muscle cells as cellular constituents. Creating anisotropic gelatin-elastin fibers will enhance the strength and toughness of the fibers and resultant constructs. It will resemble the vagina more closely, primarily when replicating the alignment of fibers and cells in longitudinal and circular directions. Advanced geometries, including replicating the tubular architecture of the vagina, will provide a more accurate testing platform, especially when incorporating geometrical and structural changes from the distal to proximal ends like the native tissue. Finally, given that vaginal smooth muscle plays critical roles in vaginal contractility and function, incorporating additional cell types like vaginal smooth muscle cells will increase the fidelity and accuracy of the model.

We recognize this system has some limitations. We will work to address these limitations in the next-generation model system but acknowledge them here. We used only one cell source for this work: primary human vaginal epithelial cells. We chose these cells as they were the only commercially available vaginal cell source at the time of this study and were previously used in a recent microfluidic vaginal model^39^. Tissue engineering techniques will enhance model complexity by creating co-cultures with additional relevant cell types. We also recognize that culturing the cells on the construct’s surface may be more representative than encapsulating the epithelial cells. We chose to encapsulate cells to ensure that the cells interacted with the entirety of the construct and to assess biocompatibility within the construct. As we incorporate additional cell types, we will culture the epithelial cells on top of the constructs using our previously established techniques to stratify cells on GelMA hydrogels^32^.

In a 2022 systematic review, Francés-Herrero et al. identified studies that used bioengineering strategies for studying the vagina, including scaffold-free techniques, decellularized ECM and polymer scaffolds, and bioprinting^40^. Some human vaginal models used scaffold-free approaches and modeled the human vaginal mucosa^41^ and human vaginal epithelium^42,43^. Scaffold-free approaches employ the self-organizing capabilities of cells to create constructs; however, their biophysical properties, such as stiffness, cannot be readily tuned. Some of the identified decellularized ECM and polymer scaffolds and bioprinted constructs studied the human vagina using decellularized porcine small intestinal submucosa, acellular human cadaver dermal matrix, collagen, alginate and chitosan, cellulose, poly-lactic acid (PLA), and decellularized porcine vagina^40^. We identified an additional article published in 2022 that used a microfluidic approach to culture vaginal cells with constituents of the vaginal microbiome^39^.

We believe our model is the first to combine electrospun fibers with hydrogels into fiber-reinforced hydrogels for vaginal tissue engineering. We believe our composites are advantageous compared to the biomaterials mentioned above used for vaginal constructs for many reasons. First, gelatin is chemically similar to collagen, inexpensive, and biocompatible.^25^ Second, chemical modification of gelatin to GelMA renders it stable under physiological temperatures, homogenous in structure, and able to be mechanically tuned. The resultant hydrogels are also viscoelastic in nature^25,44,45^. Third, chemical polymerization techniques using UV light and a photoinitiator allow for fast (seconds to minutes) polymerization. Fourth, infiltrating electrospun fibers with hydrogels increases load-bearing capacity and confers greater strength and toughness compared to homogeneous hydrogels, which tend to be brittle. Finally, our non-toxic electrospinning technique using acetic acid to spin gelatin-elastin fiber blends represents critical components of the vaginal ECM.

Basic science information on vaginal biomechanics can provide targeted ways to inform additional applications, including childbirth injury, drug delivery, and neovagina creation. Many gynecologic exposures can result in injuries to the vagina, especially childbirth. During childbirth, the vagina stretches significantly to almost five times its diameter to accommodate the delivery of the fetus^20,46^. A 9 cm fetal head experiences approximately 16 N of force at rest from the uterus, 54 N during a uterine contraction, and 120 N during a deliberate push, which in turn exposes vaginal tissue to these stresses required to deliver the fetus^47^. Despite the ability of the vagina to partially elastically recover, vaginal birth still results in vaginal tearing and injury to the pelvic floor. The prevalence of childbirth injuries is high, with approximately 85% of women experiencing perineal lacerations (vaginal tears) during vaginal delivery ^20^.

Vaginal features (e.g., tissue structure, cellular composition, ECM components) across all length scales play critical roles in vaginal structure, function, and injury (**Fig. 9**). The underlying microstructure of the vagina is essential in dictating the tissue’s function. In particular, the cells and ECM fibers in the vagina contribute to the elastic performance and recovery of the tissue in childbirth. However, much remains unknown regarding the vaginal microstructure in general, including how the fiber-cell and cell-cell interactions influence the macro mechanical properties of the tissue. A deeper understanding of these interactions will provide critical missing information needed to reduce and prevent vaginal injuries, including perineal lacerations.

**Figure 9.**
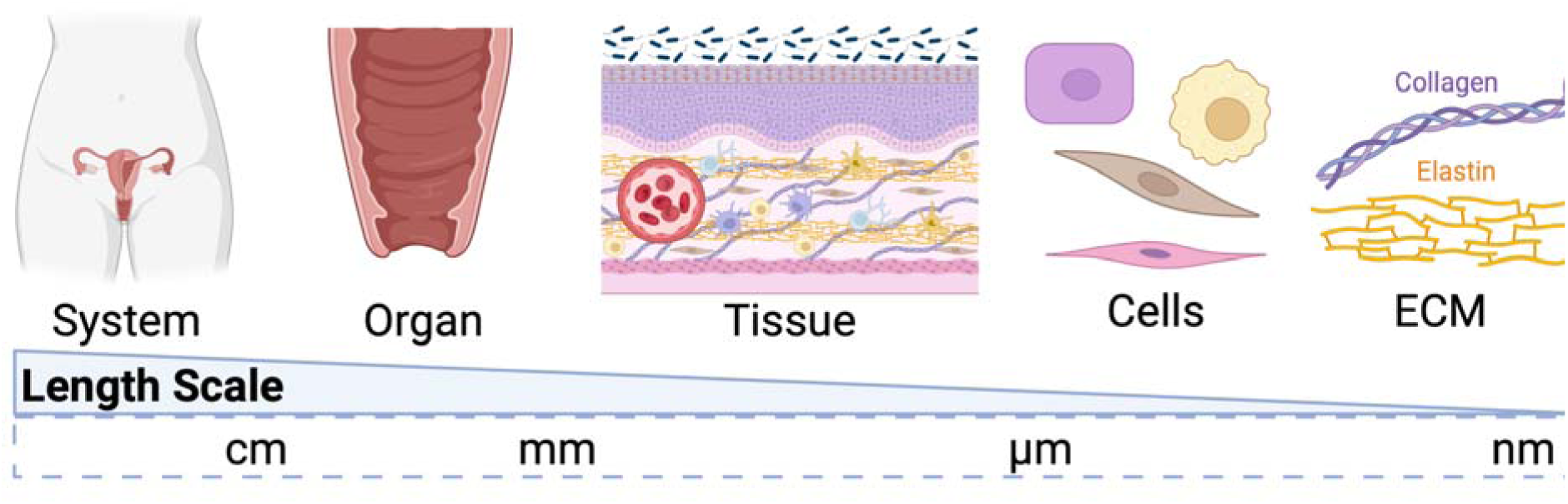
The vaginal length scale. Relative sizes of vaginal components from system to extracellular matrix (ECM) level

Electrospun fibers have been used for vaginal drug delivery, including pre-exposure prophylaxis (PrEP) drugs incorporated in polycaprolactone (PCL) and polyvinyl alcohol (PVA) electrospun fibers, antiretroviral drug-laden nanoparticles incorporated in PVA and polyvinylpyrrolidone (PVP) electrospun fibers, herpes simplex virus 2 drug incorporated in poly(lactic-co-glycolic acid) (PLGA) and poly(l-lactide-co-caprolactone) (PLCL) electrospun fibers, as well as others^48^. The advantages of electrospun fibers for vaginal drug delivery include the use of mucoadhesive polymers capable of improving the retention of vaginal drug doses and maintaining a high surface area^48^. Combining therapeutics and other signaling molecules has the potential to instruct native vaginal cells to integrate and interact with electrospun fibers.

Furthermore, the creation of a neovagina for individuals with congenital malformations, including Mayer-Rokitansky-Küster-Hauser syndrome, or individuals undergoing gender-affirming surgery is still limited, and the use of non-vaginal tissues for reconstruction is associated with complications such as graft rejection, lubrication issues, and excessive vaginal discharge^49^.

Biomechanical problems associated with neovaginal tissue, including vaginal stenosis, dyspareunia, and vaginal prolapse, can negatively affect the performance of the tissue^49^. As we continue engineering efforts for creating a tissue engineered vagina, we have the potential to target these issues and develop neovaginal constructs for vaginal reconstruction with enhanced features that reduce these complications and challenges.

We will continue to advance model complexity by achieving and adding additional benchmarks established from our predefined engineering criteria. As we continue to progress, these systems will move toward an entire tissue engineered vagina system. Advancing tissue engineered systems for modeling the vagina will provide crucial engineering tools to generate important basic science data related to vaginal tissue function.

## 5. Conclusions

The fiber-reinforced composites for vaginal tissue engineering proposed herein are an innovative platform with enhanced technical capabilities compared to existing models. This multiscale effort approached vaginal tissue engineering from material, tissue, and cell perspectives. We created novel fiber-reinforced composites containing electrospun gelatin-elastin fibers within GelMA hydrogels. We defined the material and mechanical properties of these composites and described their biocompatibility with the vaginal microenvironment. This project significantly advances progress in vaginal tissue engineering by developing novel materials for tissue engineering and developing a state-of-the-art tissue engineered vagina.

## Acknowledgments

The authors thank Dr. Huafang Li for assisting with ESEM training in the IMSE Core Facility at Washington University in St. Louis. The authors thank Dr. Adrienne K. Scott for reading this manuscript. Gregory Strout performed critical point drying through the use of the Washington University Center for Cellular Imaging (WUCCI) supported by the Washington University School of Medicine, The Children’s Discovery Institute of Washington University, and St. Louis Children’s Hospital (CDI-CORE-2015-505 and CDI-CORE-2019-813) and the Foundation for Barnes-Jewish Hospital (3770 and 4642). This work was supported by the National Institutes of Health T32 Clinical Outcomes Research Training Program in Female Lower Urinary Tract Disorders [NIH T32DK120497] (SGZ). Figures were created with Biorender.com.

We describe contributions to the manuscript using the Contributor Roles Taxonomy (CRediT):

*Conceptualization-SGZ, SS, JLL, MLO Methodology-SGZ, MLO*

*Formal Analysis-SGZ, SSK, AH*

*Investigation-SGZ (GelMA synthesis, ESEM imaging, cell experiments), SSK (tear testing, degradation), AH (electrospinning, fiber characterization, degradation, mass swelling), AMM (electrospinning, viscosity measurements)*

*Resources-SGZ, MLO*

*Data Curation-SGZ*

*Writing: Original Draft-SGZ*

*Writing: Review and Editing-SGZ, SSK, AH, AMM, SS, JLL, MLO*

*Visualization-SGZ*

*Supervision-MLO, SS, JLL Project Administration-MLO*

*Funding Acquisition-SGZ, MLO*

## Data Availability Statement

The datasets used and analyzed during the current study are available from the corresponding authors upon reasonable request.

