## Supplemental Information for "Vaginal Tissue Engineering via Gelatin-Elastin Fiber-Reinforced Hydrogels"

```
% Created by Annie Mascot, WashU, April 2023
% Calculating viscosity of a fluid from a Cannon-Fenske Viscometer



% input temperature of sample and time
temp = 23;
t = [36.49, 250.21, 411.95, 831.6, 3817.67, 3231.33];
sample = categorical({'Water', 'Elastin', '1:2', '1:1', '2:1','Gelatin'});

% obtained calibration constants and plot
p1 = [40, 0.02803];
p2 = [100, 0.02811];
x = [0:1:150];
m = (p2(2)-p1(2))/(p2(1)-p1(1));
y = m*(x-p1(1))+p1(2);
n = find(x==temp);
c = y(n);
% plot(x, y,'LineWidth',1)
% grid on
% hold on
% plot (n, c, '.k','MarkerSize', 12)
% xlabel('Temperature [¬∞C]')
% ylabel('Calibration Constant')
% xticks(0:10:x(end))
% title('Calibration Constant for Canon-Fenske Viscometer')


% calculate viscosity
for i=1:length(t)
v(i) = c*t(i); %[cst]
end

bar(v)
set(gca,'xticklabel',sample)
ylabel('Viscosity [cts]')
xlabel('Sample Type')
title('Viscosity of Polymer Solutions')
legend('Ratios Display Gelatin:Elastin. All samples are 10% [w/v]')

vdisp = num2str(v);

disp(['v = ', vdisp, ' centistokes'])
```

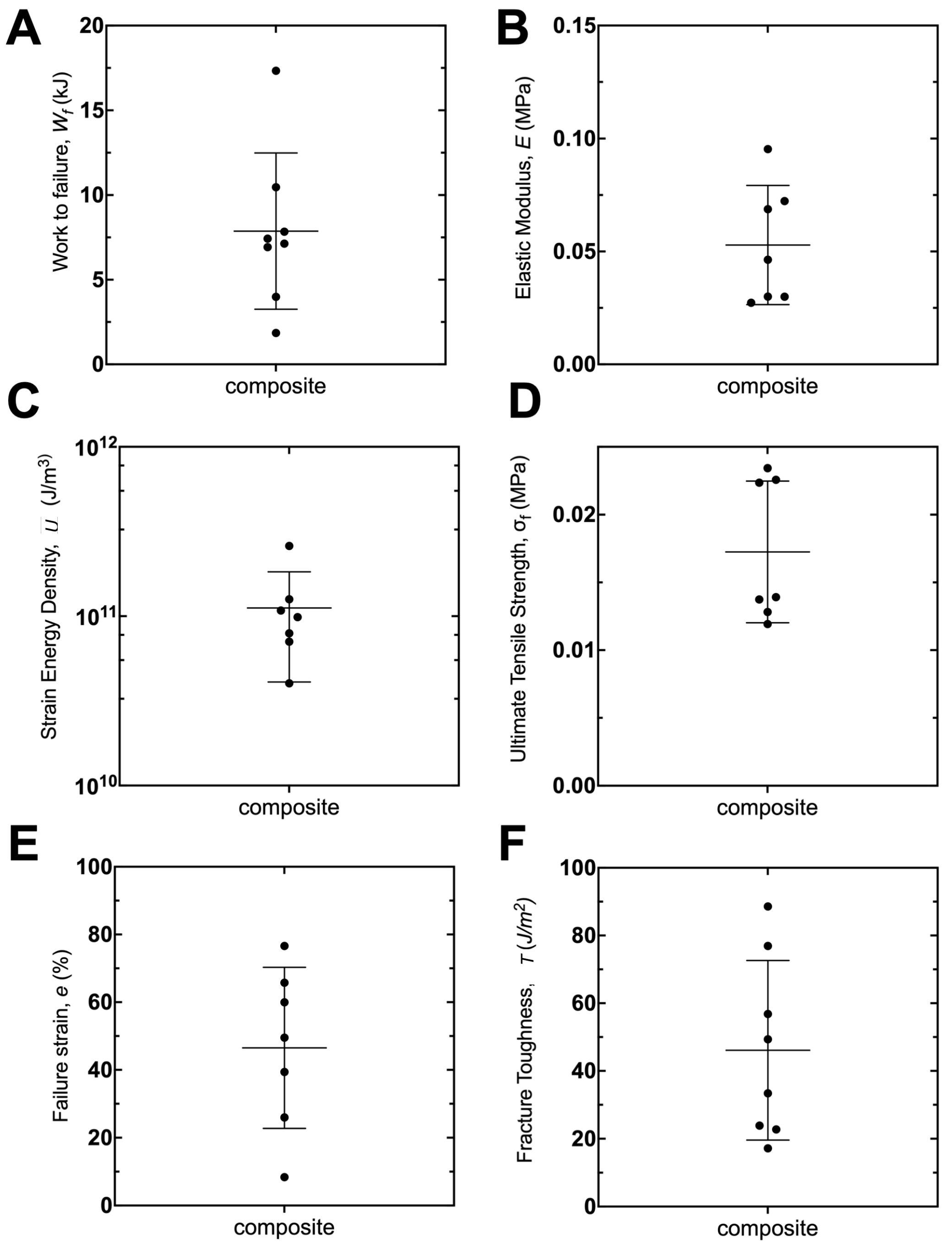


**Figure S1.** Tear testing results for fiber-reinforced composites. **A.** Work to failure. **B.** Elastic modulus. **C.** Strain energy density. **D.** Ultimate tensile strength. **E.** Failure strain. **F.** Fracture toughness. Data are presented as mean ± standard deviation.

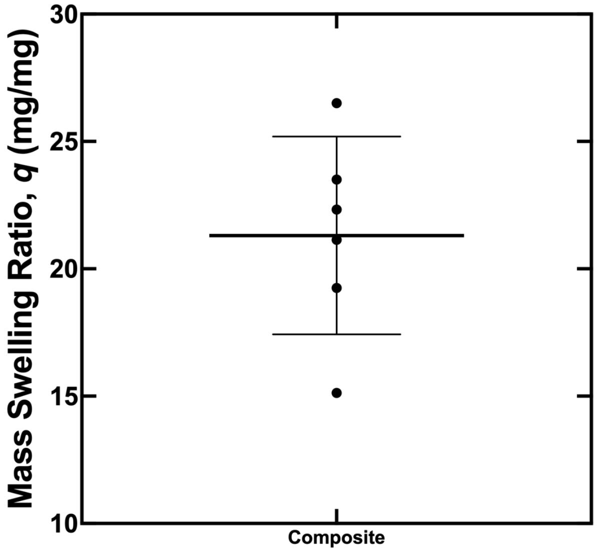


**Figure S2.** Mass swelling ratio of the composites. Data presented as mean ± standard deviation.
